# Microstructure-Based Nuclear Lamina Constitutive Model

**DOI:** 10.1101/2023.07.24.550440

**Authors:** Nima Mostafazadeh, Zhangli Peng

## Abstract

The nuclear lamina is widely recognized as the most crucial component in providing mechanical stability to the nucleus. However, it is still a significant challenge to model the mechanics of this multilayered protein network. We developed a constitutive model of the nuclear lamina network based on its microstructure, which accounts for the deformation phases at the dimer level, as well as the orientational arrangement and density of lamin filaments. Instead of relying on homology modeling in the previous studies, we conducted molecular simulations to predict the force-extension response of a highly accurate lamin dimer structure obtained through X-ray diffraction crystallography experimentation. Furthermore, we devised a semi-flexible worm-like chain extension-force model of lamin dimer as a substitute, incorporating phases of initial stretching, uncoiling of the dimer coiled-coil, and transition of secondary structures. Subsequently, we developed a 2D network continuum model for the nuclear lamina by using our extension-force lamin dimer model and derived stress resultants. By comparing with experimentally measured lamina modulus, we found that the lamina network has sharp initial strain-hardening behavior. This also enabled us to carry out finite element simulations of the entire nucleus with an accurate microstructure-based nuclear lamina model. Finally, we conducted simulations of transendothelial transmigration of a nucleus and investigated the impact of varying network density and uncoiling constants on the critical force required for successful transmigration. The model allows us to incorporate the microstructure characteristics of the nuclear lamina into the nucleus model, thereby gaining insights into how laminopathies and mutations affect nuclear mechanics.

## 1. Introduction

The nuclear lamina is a random fibrous network of lamin protein filaments that underlies the inner nuclear membrane and interfaces with chromatin in most eukaryotic cell nuclei [1]. It has long been studied for its key role in the mechanical stability, development, and transcriptional activities of the cell [2]. The constituents of the nuclear lamina are lamin filaments, which are tetrameric assemblies of polymerized lamin isomers [3] including lamin A (lamin A and C) and lamin B types (lamin B1 and B2), which encoded by the LMNA and LMNB gene groups, respectively. Furthermore, studies have shown that lamin isomers form separate meshworks with unique functionalities and organizational structures [4]. Research suggests that the lamin A network, in particular, plays a critical role in maintaining the mechanical integrity of the nucleus during deformation events throughout its lifespan [5].

The nuclear lamina regulates and influences numerous critical biological events in cells. Therefore, structural malfunction of the nuclear lamina can lead to or facilitate severe conditions that pose significant risks, such as cancer and viral infections [6]. Laminopathies, which are attributed to such failures in the nuclear lamina, encompasses various types of muscular and skeletal dystrophies, dilated cardiomyopathy, neuropathies, and other related conditions [7]. Additionally, it includes a rare yet complex premature aging disorder known as Hutchinson-Gilford progeria syndrome (HGPS) [7]. Certain conditions are the result of disrupted cellular proteostasis activity and the localization of mutant lamin isomers in the nuclear lamina [8]. Mutations in lamin A can alter its network-level assembly and behavior, resulting in laminopathic and cancerous phenotypes [9]. In the case of HGPS, a single point mutation in the LMNA gene leads to the transcription of progerin, a protein that replaces lamin A in the nuclear lamina, and results in the formation of dysfunctional networks [10]. In addition, It has been observed that lamin A is either upregulated or downregulated in various types of cancer [11], contributing to the deterioration of nuclear biophysics and mechanically over-softened or over-stiffened response. These deviations significantly impact cell vitality and activity during transmigration through confined spaces [12-16], endothelial gaps [17] and the bloodstream [18]. The significance of these physical phenomena has sparked an interest in utilizing mathematical models and simulations to better comprehend nuclear deformation during related processes.

Numerous efforts have been made to develop models that depict the mechanical behavior of the nuclear lamina [19]. These models have been employed in standalone simulations as well as in simulations of the entire nucleus and they encompass a wide range of approaches, including continuum linear or nonlinear shells and coarse-grained polymer models. The parameters of these models have been carefully calibrated to experimental data, ensuring their fidelity to real-world observations. Research indicates that the nuclear lamina is a hierarchical, multilayered protein network characterized by its dynamic and composite assembly. Laminopathies affect cross-sectional bonds and the local concentration of lamin is directly influenced by both its nucleus transcriptive health and the mechanical deformations it experiences [20]. Consequently, mathematical models that are suitable for geometrically or chemically homogeneous structures may lack the flexibility required to investigate the variable components of the lamin network microstructure, which assertedly, are responsible for the physiological basis of mechanical deviations observed *in vivo*. As a result, subsequent attempts have been made to model the implication of the nuclear lamina as a meshwork.

Qin and Buehler developed a multi-linear relationship fitted to the *in silico* force-extension response of a lamin dimer structure generated from homology modeling and iteratively simulated a coarse-grained rectangular arrangement of filaments to capture the strain-stiffening and rupture point of the network [21]. They later simulated the same filaments in an electron-tomography image-based network assembly [22]. These molecular dynamic simulations, aside from being time costly, present challenges when it comes to integrating with other cellular components for larger-scale simulations. A preferable approach is to develop a continuum model that takes into account the hierarchical mechanics and arrangement of the nuclear lamina and is responsive to its physical characteristics.

Cao et al. [23] modeled tumor cell migration in microfluidics and included the nuclear envelope as an empirical hyperelastic material and chromatin as an empirical poroelastic material. To consider the external force applied on the nucleus, they applied a chemomechanical model based on actin-fiber contraction. Recently, Arefi et al. proposed a biomechanical model for the extravasation of a tumor cell (TC) through the endothelium of a blood vessel [24]. They assume that the TC extends a protrusion between adjacent endothelial cells (ECs) that adheres to the basement membrane via focal adhesions (FAs). The pulling force in the TC invadopodia is modeled by adopting models by Deshpande et al. [25-27] for the SFs and the FAs, both controlled by biochemical signaling with a mechanical feedback. However, the nucleus was still modeled as a homogeneous solid using empirical constitutive law, and the microstructures of the nuclear envelope and chromatins were completely ignored.

In this study, we elucidate the importance of the experimentally derived crystal structure used for our molecular dynamics (MD) uniaxial stretching simulation. We propose a mathematical force-extension model for the extension of a coiled-coil structure as a function of the applied pulling force, which bridges the low-force and high-force regimes in MD force-displacement data using a two-state kinetic model. Subsequently, we develop a constitutive model that integrates the coiled-coil force-extension model into a 2D continuous network, enabling its implementation in finite element simulations. Additionally, to estimate the initial shear modulus and the network density, we utilize the developed coiled-coil model and Cryogenic Electron Tomography (Cryo-ET) imaging analysis, respectively. Finally, we compare the effect of various intrinsic parameters of the model in the nuclear lamina mechanics and the computed critical force in the finite element simulation of nucleus transmigration through endothelial pores.

## 2. Mathematical Formulation

### 2.1. Lamin Dimer Structure

To investigate the mechanical behavior of the full-atom dimer structure, we conducted a uniaxial stretching molecular dynamics (MD) simulation by applying a constant velocity of 50 Å/ps to the head end of the structure along the direction of the helical axis and recorded the force-displacement data associated with the stretching process throughout the simulation. In contrast to the previously employed homology-based predicted structure of lamin A which has been used in past lamin mechanics-related models [21, 22], we utilized the experimentally derived crystal structure obtained by Ahn et al. They managed to determine the first 300 residues of the lamin A protein at a resolution of 3.2 Å, enabling us to elucidate a substantial portion of its three-dimensional structure (Fig. 1a) [28]. The studied domains, namely, begin with N-terminal head, followed by coil 1a, L1 linker, coil 1b, L12 linker and approximately half of the coil 2, respectively. To analyze the full dimer organization, they integrated their crystal configuration with previously reported fragments (comprising the remainder of coil 2 and the tail domain) obtained from the literature. Their findings revealed that the primary composition of a lamin dimer is predominantly comprised of coiled-coil superhelical regions. Through lateral association, dimers align their coil 1b and coil 2 regions in an anti-parallel manner to form tetramers. These tetramers then undergo polymerization to generate protofilaments, which are considered the most stable fibers in the nuclear lamina network at the highest level (Fig. 1b) [3]. The protofilaments overlap in apparently arbitrary alignments with respect to each other to form the nuclear lamina meshwork, as depicted in the Cryo-ET image shown in Fig. 1c. In Fig. 1d, a schematic representation of a representative volume element (RVE) of the nuclear lamina showcases how a sub-section of how these alignments appear.

**Fig. 1.**
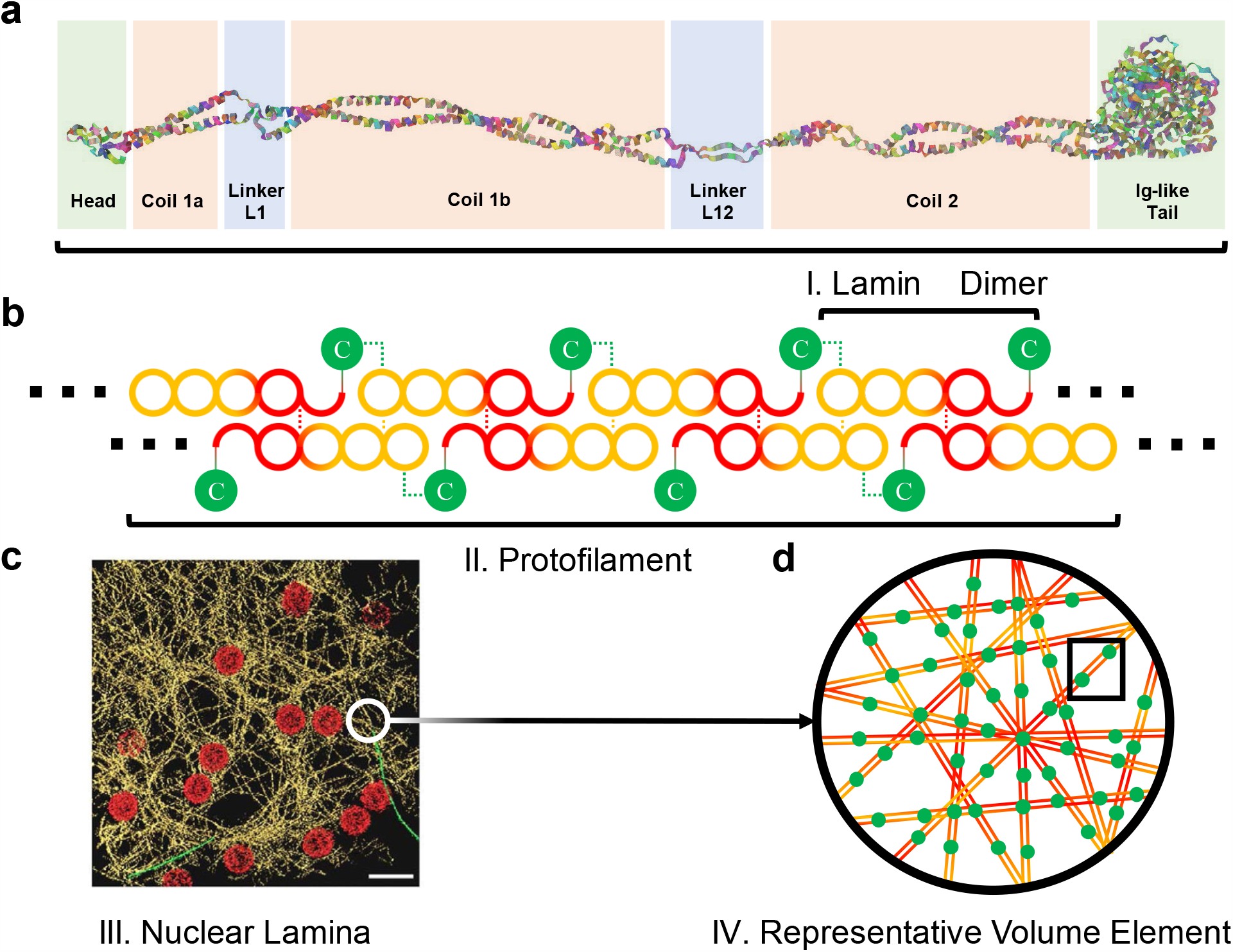
Nuclear lamina hierarchical structure. (a) The crystal structure of the lamin dimer is depicted by ribbon model, accentuating superhelical coil domains in red, linkers in blue, and head and tail residues enclosed within green boxes. This visualization clearly underscores that the structural properties of lamin are predominantly regulated by its coiledcoil motifs. (b) The high-order protofilaments, formed through a tetrameric arrangement of dimers in an anti-parallel head-to-tail configuration, are schematically demonstrated, with the C-terminal immunoglobin-like domain denoted by green circles. (c) The cryo-ET image of the nuclear lamina meshwork reveals irregularly distributed and randomly aligned lamin protofilaments, highlighted in gold, alongside nuclear pore complexes (NPCs) represented by red circles. (d) The crystal structure of lamin dimers, their lateral and axial assembly pattern, and their interpreted organization within a network form the basis for the introduction of microstructure-based model in this research describing the constitutive mechanics of the nuclear lamina.

### 2.2. Lamin Dimer Extension Model

We developed a kinetic-based two-state formulation for a semiflexible filament, specifically tailored to model the extension behavior of the coiled-coil proteins such as lamin dimer in response to stretching forces. We utilized the Blundell and Terentjev (BT) extension-force equation in our modeling approach to capture the pre-uncoiling stiffening behavior of the coiled-coil [29] (Fig. 2a-I):

**Fig. 2.**
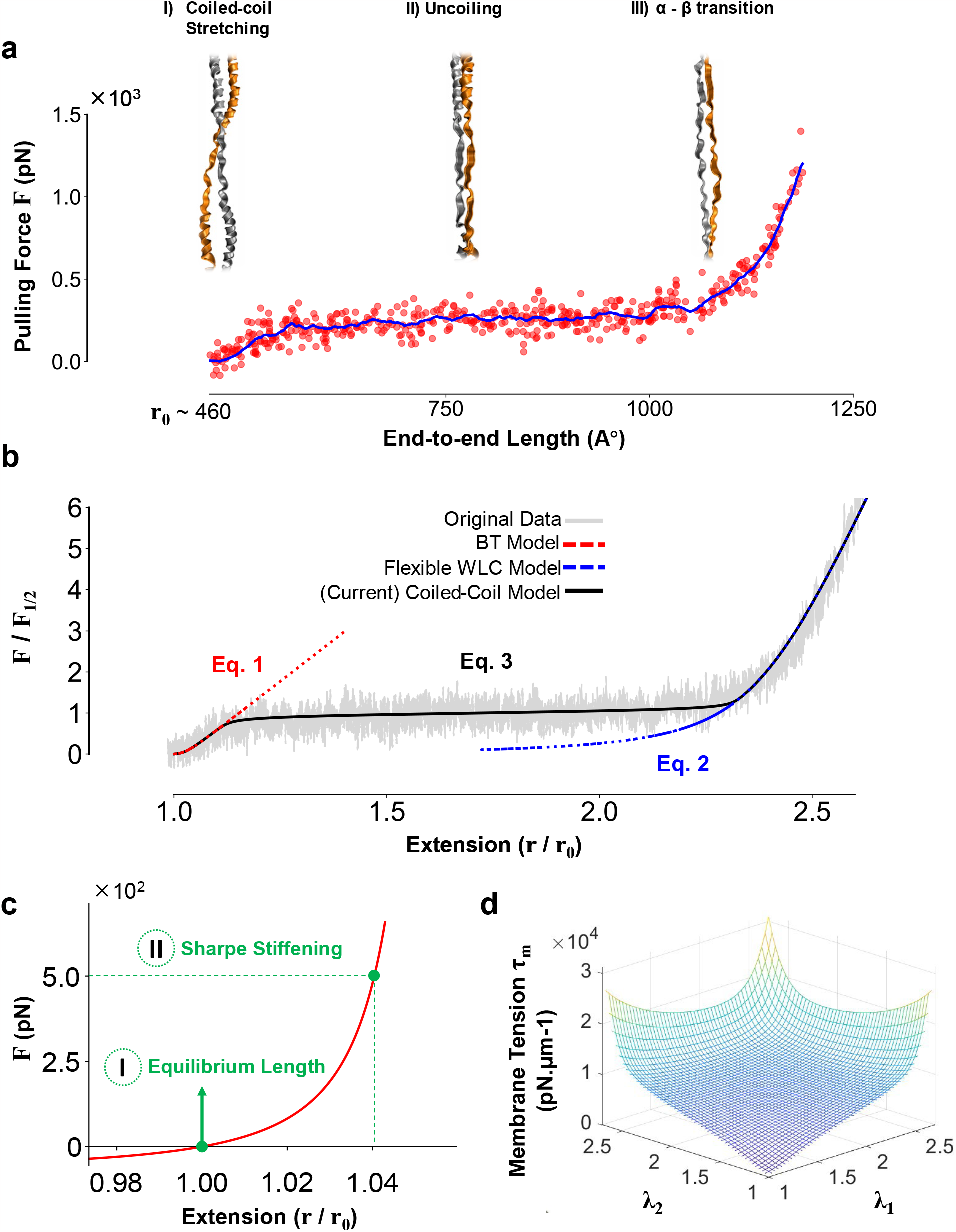
Hierarchical modeling of lamin dimer MD force-extension curve and the nuclear lamina 2D network. (a) The figure illustrates lamin dimer conformations during different deformation regimes. Initially, the dimer experiences stiffening as the coiled-coil region is stretched, overcoming conformational entropy. Subsequently, uncoiling occurs at nearly constant force, followed by a final stiffening phase caused by the transition from alpha helices to beta sheets. The initial and final stiffening regimes may signify a reinforced and stable natural state, along with constrained softening behavior of the network. (b) Semiflexible coiled-coil model (Eq. 3) is fitted to the MD data. Given *r*_*0*_ = 460 Å, the parameters for Eq. 1 were determined by fitting to the initial stiffening rise, resulting in *L*_*c*_*/r*_*0*_ ∼ 1.06, *P*_*c*_ ∼ 160 nm, *β*_*0,c*_ ∼ 5×10^4^ pN/μm. For Eq. 2, the fitting to the final stiffening rise yielded estimated values of *L*_*u*_*/r*_*0*_ ∼ 2.5, *P*_*u*_ ∼ 4 A°, *β*_*0,u*_ ∼ 1.8 × 10^3^. Subsequently, based on the previous estimates, *ΔΔx*^*\**^ = 3 A°, and *F*_*1/2*_ = 290 pN were approximated for Eq. 3. (c) The BT model incorporates two important features: 1) it inherently considers the equilibrium length of the filament, and 2) it offers steeper stiffening behavior, resulting in a more stable shear modulus in early deforming stages. (d) Membrane tension of the nuclear lamina network (Eq. 14) with respect to principal stretches *λ*_*1*_ and *λ*_*2*._

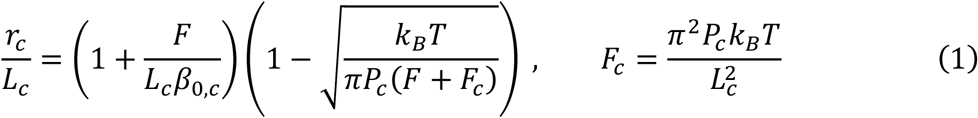

where subscript “*c*” is used to denote that the parameters for dimer end-to-end length (*r*_*c*_), contour length (*L*_*c*_), persistence length (*P*_*c*_) and extensibility constant (*β*_*0,c*_) are specifically defined for the coiled filament. *F* is the applied force, *k*_*B*_ and *T*are Boltzmann constant and temperature. To account for the behavior of highly flexible uncoiled filaments in less entropic extended dimer, we employed Petrosyan interpolation of the highly extended limit of wormlike chain (WLC) model for flexible protein. This model effectively explained the observed trend in the high-force regime [30] (Fig. 2a-III):

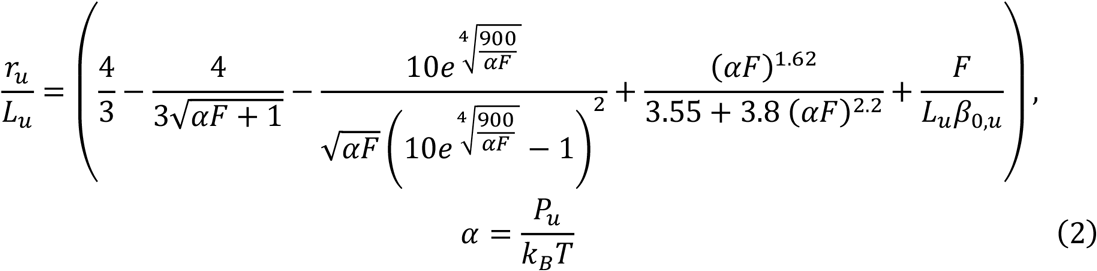

where subscript *u* in here indicates that variables *P*_*u*_, *L*_*u*_, *β*_*0,c*_ are persistence length, contour length and extensibility constant defined for uncoiled dimer. A two-state transition kinetic model is used to introduce the fraction of uncoiled over coiled length (*ϕ*_*u*_) under equilibrium to integrate *r*_*c*_*/L*_*c*_ from Eq. 1 and *r*_*u*_*/L*_*u*_ from Eq. 2 and model the uncoiling process (Fig. 2a-II):

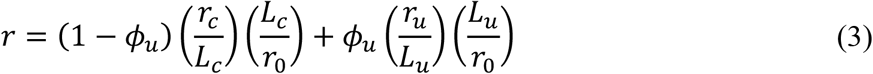

where *r*_*0*_ is defined as the initial length of the dimer and equals to 460 Å. At equilibrium, the coiling and uncoiling rates are equal i.e., *l*_*c*_*k*_*c→u*_*=l*_*u*_*k*_*u→c*_, where *l*_*c*_ and *l*_*u*_ are length of coiled and uncoiled segments (*r=l*_*c*_*+l*_*u*_) and *k*_*c→u*_ and *k*_*u→c*_ are constants for respective transitions rates. The relationship between *r*_*c*_and *l*_*c*_is expressed as *l*_*c*_*=*(1−*ϕ*_*u*_)*r*_*c*_, and *r*_*u*_ is related to *l*_*u*_ through *l*_*u*_*=ϕ*_*u*_*r*_*u*_. Here, *r*_*c*_ and *r*_*u*_ are based on whether the entire dimer follows the BT model for coiled filaments or the Petrosyan interpolation for uncoiled filaments, respectively. Subsequently, one can express *ϕ*_*u*_ using Arrhenius equation in terms of temperature and coiling and uncoiling Gibbs free energies (*ΔG*_*c→u*_ and *ΔG*_*u→c*_):

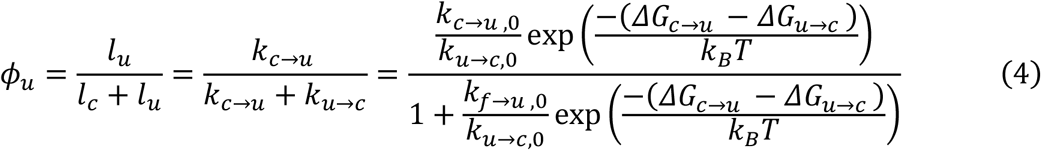

The expression can be simplified by relating the difference in transition Gibbs energies to applied force by introducing uncoiling lengths *ΔΔx*^*\**^, similar to other related studies [31]:

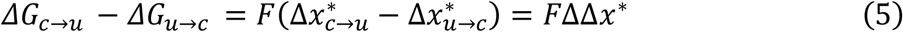

By solving *ϕ*_*u*_ *= 0*.*5* for 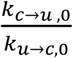 in terms of *ΔΔx*^*\**^ and uncoiling force *F*_*1/2*_, which is the force required to uncoiled half of the coiled-coil dimer, and replacing in Eq. 4, the overall expression for *ϕ*_*u*_ will be:

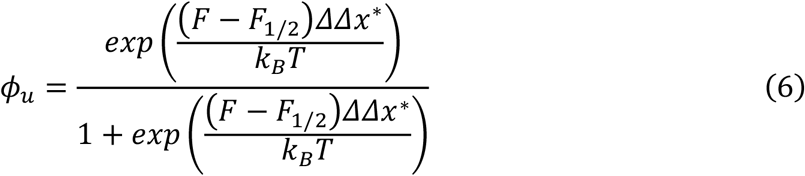

The Savitzky-Golay filter is employed to smoothen the raw force-displacement data before applying the non-linear least-squares method to fit the parameters. In order to minimize the number of independent variables in each fitting session, the data is segmented into three batches. The first batch (initial stiffening regime) is fitted to BT model (Fig. 2c) for *L*_*c*_, *P*_*c*_, and *β*_*0,c*_,while the last batch (final stiffening regime) is fitted to the Petrosyan interpolated relationship for *L*_*u*_, *P*_*u*_, and *β*_*0,u*_. Using the known coiled and uncoiled filament model parameters as a reference, the uncoiling transition constants (*ΔΔx*^*\**^ and *F*_*1/2*_) are fitted to the entire dataset, thereby quantifying all the model parameters (Fig. 2b). A summary of all model parameters is provided in Table 1.

**Table. 1.**
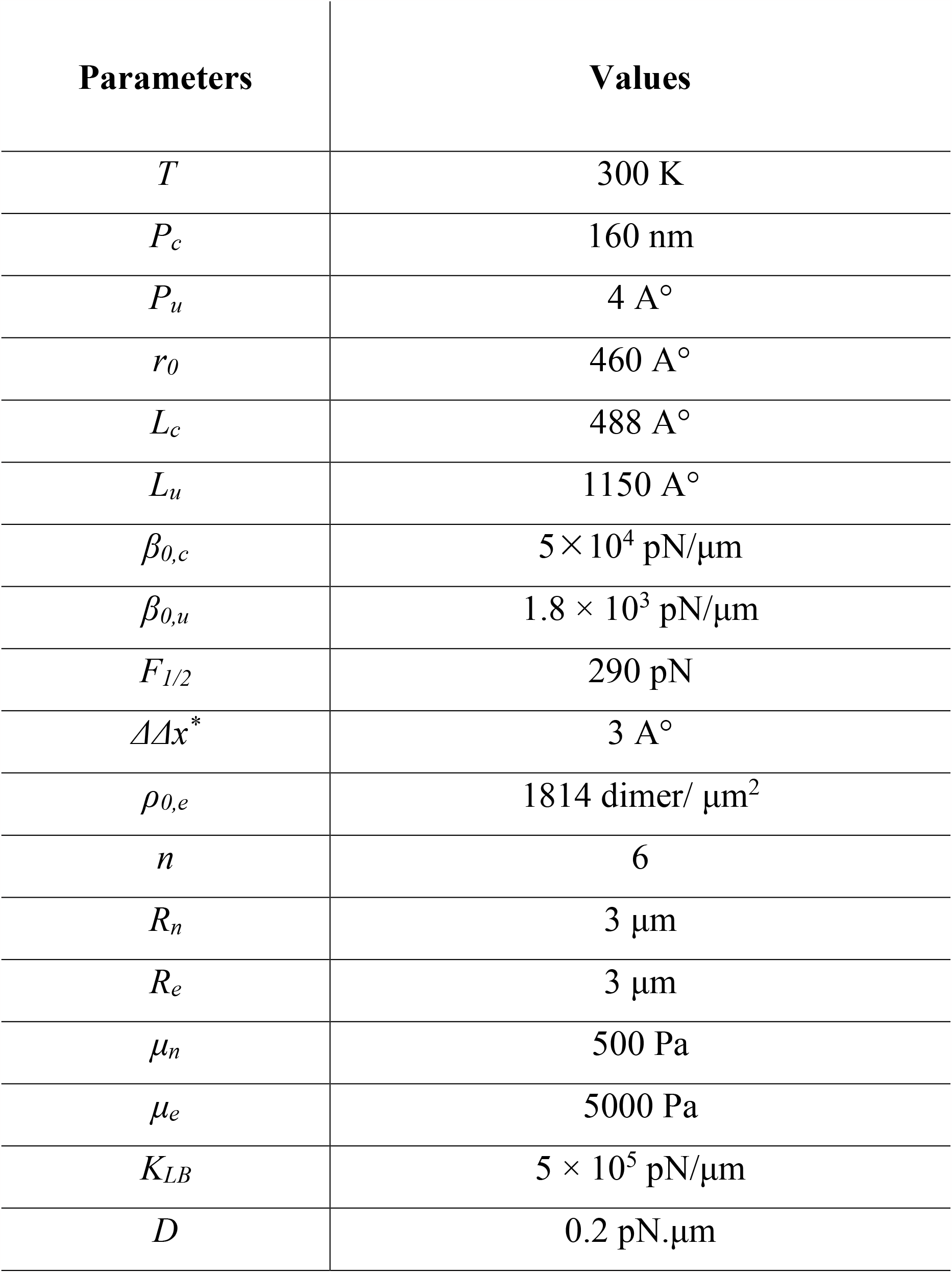
Summary of model parameters.

### 2.3. Nuclear Lamina Microstructure-Based Constitutive Model

Studies have demonstrated that the lamin A network exhibits a loose but highly dense lamin meshwork, which responds viscously to mechanical strain. On the other hand, the lamin B network forms a less dense crosslinked structure, displaying elastic behavior [20, 32]. Consequently, in deformations occurring over small timescales, the network’s topology is not anticipated to undergo significant reorganization, and sliding effects are not expected to dominate even at considerably high extensions of lamin filaments [22]. Given the above considerations, it is reasonable to assume that the network deforms affinely under lower time scale biologically relevant deformations.

Under the affine assumption, the extension *x* of lamin dimer, oriented at an angle *θ* with respect to local coordinates, is related to principal stretch ratios (*λ*_*1*_ and *λ*_*2*_) of deforming plane that contains it through following relation:

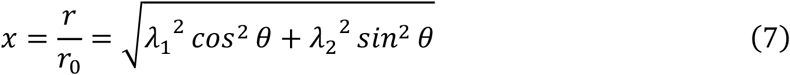

We deduced the structural properties of our network model, including the total number of dimers, using Cryo-ET image of nuclear lamina as presented in the work by Turgay et al. [33] and incorporated their analysis of filament lengths in our research. We selected the image they provided (Fig. 1c) as a RVE for the entire nuclear lamina network and further deconstructed the structure into rigidly attached dimers. Thus, the total network energy for an RVE of known dimensions can be expressed as follows:

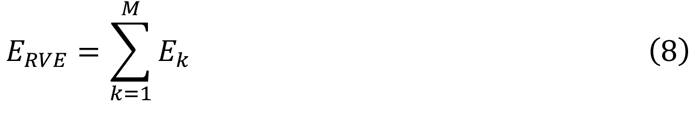

where *M* is the total number of dimers in the RVE. To calculate the total energy, we organized the dimers into multiple sets based on specific characteristic orientations. Under the assumption of uniform randomness in filament orientation in vivo, we assigned an equal number of dimers to each set:

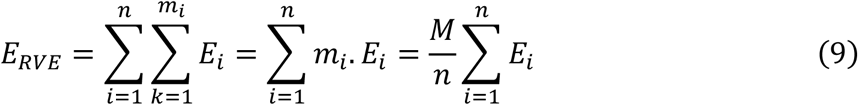

where *n* is the number of characteristic orientations and *m*_*i*_ is the number of dimers in the RVE with orientation *i*. The energy density function of the RVE (*W*_*RVE*_) is determined by dividing the *E*_*RVE*_ by its rested area (*A*_*0*_). By establishing a relationship with the initial network density (*ρ*_*0*_), we expressed the total number of dimers and RVE rested area as follows:

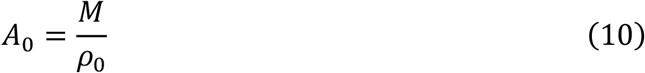

The internal energy increment in a filament (*dE*) is equivalent to the work needed to gradually stretch it (*F*.*dr*). Consequently, the stress resultants of the RVE for two principal directions have been solved analytically:

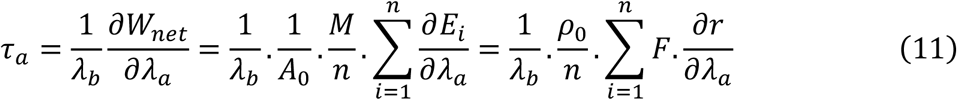

where F is the force.

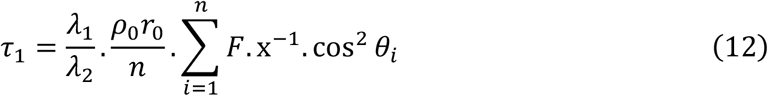

In an axisymmetric 2D scheme, *a, b ϵ*{1, 2} and *a ≠b*, therefore:

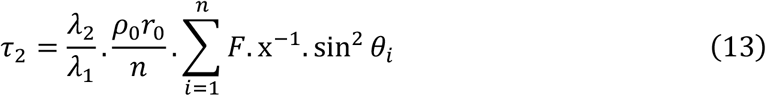

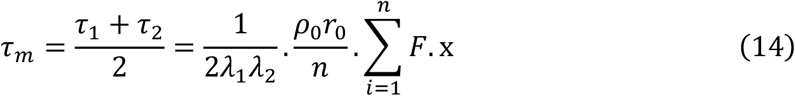

The membrane tension, shear stress resultant and shear modulus of the RVE are then derived respectively as:

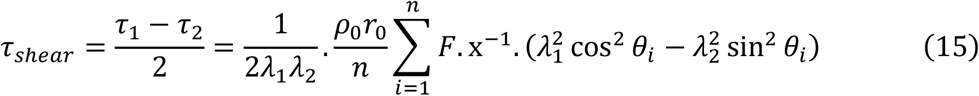

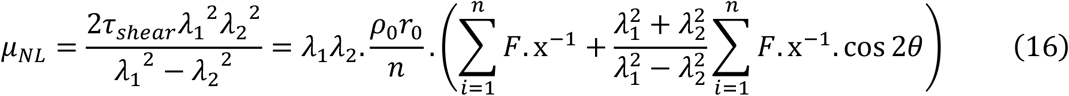

where *μ*_*NL*_ is the shear modulus of the nuclear lamina (NL).

Fig. 2d graphs the membrane tension against the principal stretch ratios *λ*_*1*_ and *λ*_*2*_, after substituting Eq. 3 in Eq. 14.

## 3. Results

### 3.1. Force-Extension Curve

The force-extension data of the lamin A dimer exhibits striking similarities to those of macromolecules with common structural characteristics, such as vimentin [34], myosin [35, 36], and double-stranded DNA [37], which can be attributed to the presence of shared features, including coiled-coils and extended helical repeats. On this account, all of these molecules exhibit similar stretching regimes, characterized by an initial rise, followed by a plateau, and concluding with a final rise (Fig. 2a). At the outset of stretching, the substantial amount of bending energy stored in coiled-coil structure transforms into mechanical work as the molecule is further stretched. The conversion continues as the entire backbone gradually aligns with the direction of the applied force. At this point, the dimer begins to uncoil, a distinctive phase of constant force deformation which is natural to coiled-coil setup [38]. Following the uncoiling of the dimer into two monomeric strands, the alpha helices undergo a secondary structure transition into beta sheets [39]. As the backbone becomes increasingly enthalpic, the required force for stretching it begins to increase. Collectively, these processes constitute the final rise phase [35]. Sapra et al. captured similar deformation levels by performing atomic force microscopy (AFM) indentation on single lamin filaments in situ, confirming the significance of coiled-coil compartment in lamin dimer [22].

Flexibility of protein conformations are compared by the ratio of their contour length (*L*) and persistence length (*P*), which under Brownian fluctuations is related to bending rigidity of the chain (*B*) by 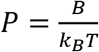 where *k*_*B*_ is the Boltzmann constant and *T* is absolute temperature. [40]. The comparable contour and persistence length in undeformed lamin dimer guarantees just enough bending rigidity to conserve rod-like structure [41]. Under a largely acceptable range, this state is known as semi-flexibility [42]. Since beta-sheets are more flexible than alpha-helices [43, 44], as the protein unfolds, the persistence length of filament falls drastically behind its contour length and the whole protein destines to act flexible. WLC model fits suitable to describe mechanics of both semi-flexible and flexible filaments with finite bending rigidity. The model is a continuum approach to evaluate bending energy of a filament based on local curvature under Brownian fluctuations, that is:

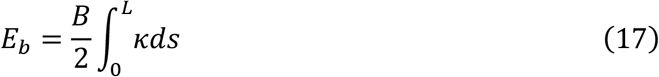

where *κ* is local curvature at arc point *s*. Many solutions to the WLC integral focus on the limiting behavior of highly extended polymers, making them inadequate for describing the initial state of a semiflexible filament. Blundell and Terentjev successfully derived a physics-based free energy which relates to the polymer extension over a wide range, leading to an insightful force expression (BT model) for semi-flexible polymers.

By fitting the BT model for semiflexible filaments to the initial stiffening regime (Fig. 2b), rather than relying on the linear behavior typically described in the literature [36, 38], we obtained a persistence length (*P*_*c*_ *≈* 160 nm) that falls within the range reported in existing literature [33]. In contrast to the flexible high-force limit solutions of the WLC model, the BT force-extension model incorporates a natural equilibrium length for the filament (set to *r*_*0*_ = 460 A° for lamin A dimer) and exhibits a comparatively sharper stiffening response, which in turn provides network a higher primary shear modulus (Fig. 2c). Notably, the fitted persistence length is larger than the contour length (*L*_*c*_ *≈* 50 nm), indicating the semi-flexible nature of this phase. The analysis of the fitted persistence lengths reveals that the entropic coiled dimer exhibits a substantially higher persistence length compared to the uncoiled extended protein *P*_*u*_ *≈* 4 A°, which also aligns with the persistence length of a random coil [45]. The extensibility constant carries distinct meanings for the coiled and uncoiled stiffening regimes. For the former (*β*_*0,c*_ *≈* 50,000 pN/um), it may represent the extension of random coil regions, whereas for the latter (*β*_*0,u*_ *≈* 18,000 pN/um), it characterizes the stretching of hydrogen bonds in beta-sheets.

### 3.2. Orientation

Fig. 3 illustrates the influence of the number of characteristic orientations *n* on network membrane tension *τ*_*m*_ (Eq. 14) and shear modulus *μ*_*NL*_ (Eq. 16) for uniaxial straining scenario in principal directions. Fig. 3a serves as both a legend for the graphs and showcase of the symbolic representations for different values of n. In all the cases presented, except for the one with *n* = 3, a symmetric structure is observed. This asymmetry leads to a corresponding asymmetric response between the principal stress resultants *τ*_*1*_ and *τ*_*2*_, as resurfaced in the different membrane tension and shear modulus trends between two uniaxial straining cases (Fig. 3b-e). While the graphs display a distinctive network response for a smaller number of characteristic orientations, the effect rapidly converges as more orientations are included, bringing the network closer to a continuum plane. Super-resolution microscopy [4, 20] shows that lamin filaments form polygon faces with mostly 3 or 4 edges and junctions with 3 or more connectivity, thus, due to structural randomness, we fixed *n* to be equal to 3 for consistency. Dahl et al. [43] measured the elastic modulus of the nuclear lamina as 25 mN/m, Comparing to this value, our model prediction (Fig. 3d-e) shows that the initial shear modulus might be one order lower (∼ 2mN/m), but the shear modulus rises to about 25 mN/m with a slight deformation of 5% strain. This sharp initial strain-hardening behavior can be explained by the semi-flexible BT model. As shown in Fig. 2c, the equilibrium position (zero force) in BT model is about *x* = 0.83, which is close to the contour length limit so that its slope rises fast even if there is only 5% strain. For a semi-flexible polymer with a persistence length larger than its contour length, such as the lamin dimer studied here, the corresponding BT model predicts a sharp strain-hardening behavior in the beginning of the deformation, Since Dahl et al. [43] applied micropipette aspiration to measure the lamina modulus and its strain can be more than 5%, the modulus they obtained is an apparent modulus averaged over a range of deformation, as shown by our model predictions.

**Fig. 3.**
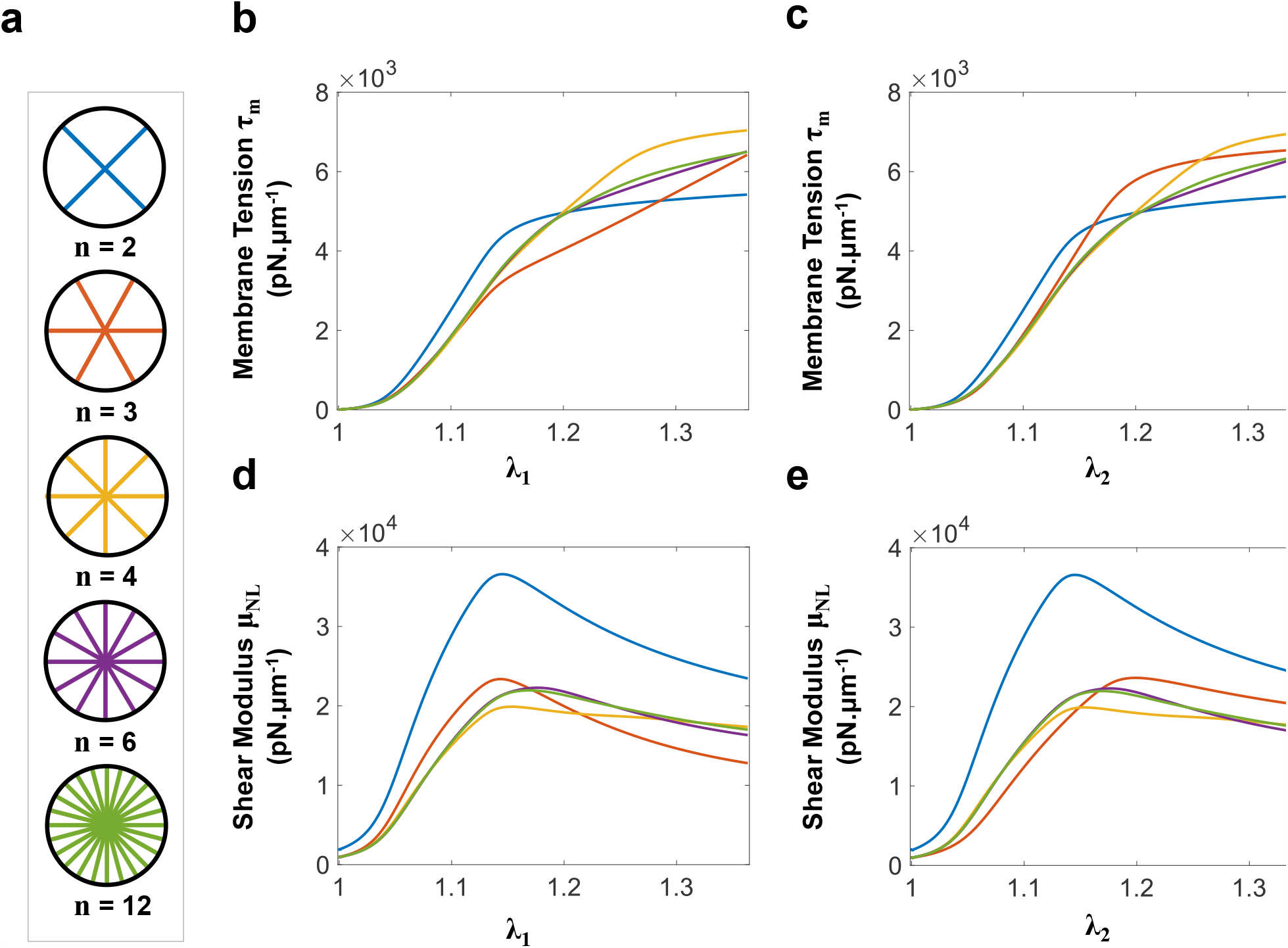
Analyzing the impact of the number of representative orientations (*n*) on membrane tension (Eq. 14) and shear modulus (Eq. 16) during uniaxial stretching. (a) The figure depicts the symbolic representation of filaments within an RVE for different values of n, where *n ϵ*{2, 3, 4, 6, 12}. For simulating purposes, *n* = 6 was selected. (b) Membrane tension (Eq. 14) is graphed with respect to *λ*_*1*_ with *λ*_*2*_ = 1. (c) Membrane tension is graphed against *λ*_*2*_ while *λ*_*1*_ = 1. (d) Shear modulus (Eq. 16) is graphed with respect to *λ*_*1*_ with *λ*_*2*_ = 1. (e) Shear modulus is graphed against *λ*_*2*_ while *λ*_*1*_ = 1. It’s worth noting that both the membrane tension and shear modulus curves exhibit distinct trends during uniaxial stretching along the 1st and 2nd principal directions when *n* = 3. This asymmetry in response is expected for all odd values of *n*. Convergent behavior of both variable as a function of filament selected representative orientation is evident. Moreover, the initial stiff and entropic response observed in the semi-flexible lamin dimer contributes significantly to the mechanics of the nuclear lamina, as it offers a substantial stability buffer to tensile and shear stiffness of the network. This attribute highlights the superiority of the BT model over the wormlike chain flexible model in predicting semi-flexible behavior.

Notably, the initial rising phase of the lamin dimer, as represented by the BT model, appears to offer a potential stability buffer for the lamin network, as evident in the shear modulus response illustrated in Fig 3d-e. This suggests a unique mechanism for preserving the geometric shape of the network against typical cytoskeletal and inner-nuclear forces. Alongside the stiffening regime of the dimer, this phenomenon may indicate a permissible range for successful transmigration.

### 3.3. Estimating Initial Network Density

To estimate the density of lamin dimers in the nuclear lamina *ρ*_*0*_, we utilized the image analysis performed by Turgay et al. with Cryo-ET images of the nuclear lamina [33]. Table 2 displays the distribution of lamin filaments with different lengths, extracted from their imaging data. Given the imaged area of 1.4 μm × 1.4 μm (Fig. 1c):

**Table. 2.** Length distribution of lamin filaments from Cryo-ET image analysis [33].

| **Lamin protofilament length (nm)** | **Frequency of occurrence** |
| --- | --- |
| 200 | 10 |
| 250 | 36 |
| 320 | 59 |
| 370 | 32 |
| 430 | 34 |
| 490 | 22 |
| 550 | 12 |
| 610 | 5 |
| 670 | 4 |
| 730 | 1 |
| 790 | 1 |
| 850 | 1 |

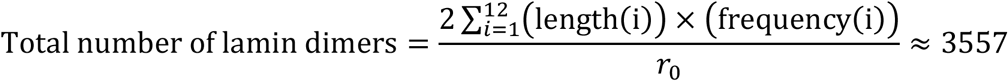

Where factor 2 is to accommodate for tetrameric cross-section of protofilaments. Consequently, the network density is estimated as:

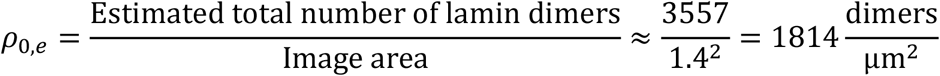

### 3.4. Finite Element Modeling

We constructed and simulated an axisymmetric finite element model of a nucleus transmigrating through an endothelial pore using Abaqus/Explicit. In Fig. 4a, the geometry used is depicted, with *R*_*n*_ representing the radius of the nucleus, *R*_*e*_ being the radius of the endothelial cell, and *δ* denoting the radius of the pore. The endothelial cell is modeled as a rigid structure; however, simulations are also conducted by assigning neo-Hookean hyperelastic properties (shear modulus *μ*_*e*_ = 5000 Pa) to endothelial cells to verify the applicability of the model in more realistic environments (Fig. 4b). The nucleus is characterized by a solid interior with neo-Hookean properties to represent the chromatin and nucleoplasm, where the shear modulus is *μ*_*n*_ = 500 Pa. Additionally, the nucleus is covered by a shell representing both the lipid bilayer and the nuclear lamina. The stiffness contribution of the lipid bilayer to the shell is incorporated by introducing an additional membrane tension based on the area modulus *K*_*LB*_ = 500,000 pN/μm and bending rigidity *D* = 0.2 pN·μm. The geometric and material values are adopted form Cao et al. [23]. We integrated the Abaqus user material definition module VUMAT to implement our model in Abaqus. A shear force is applied to the upper half of the nucleus, causing it to push through the pore. The resistance force exerted on the endothelial cell is then measured and recorded as the critical force required for successful transmigration. Fig. 4b shows the nucleus at its critical position before passing through.

**Fig. 4.**
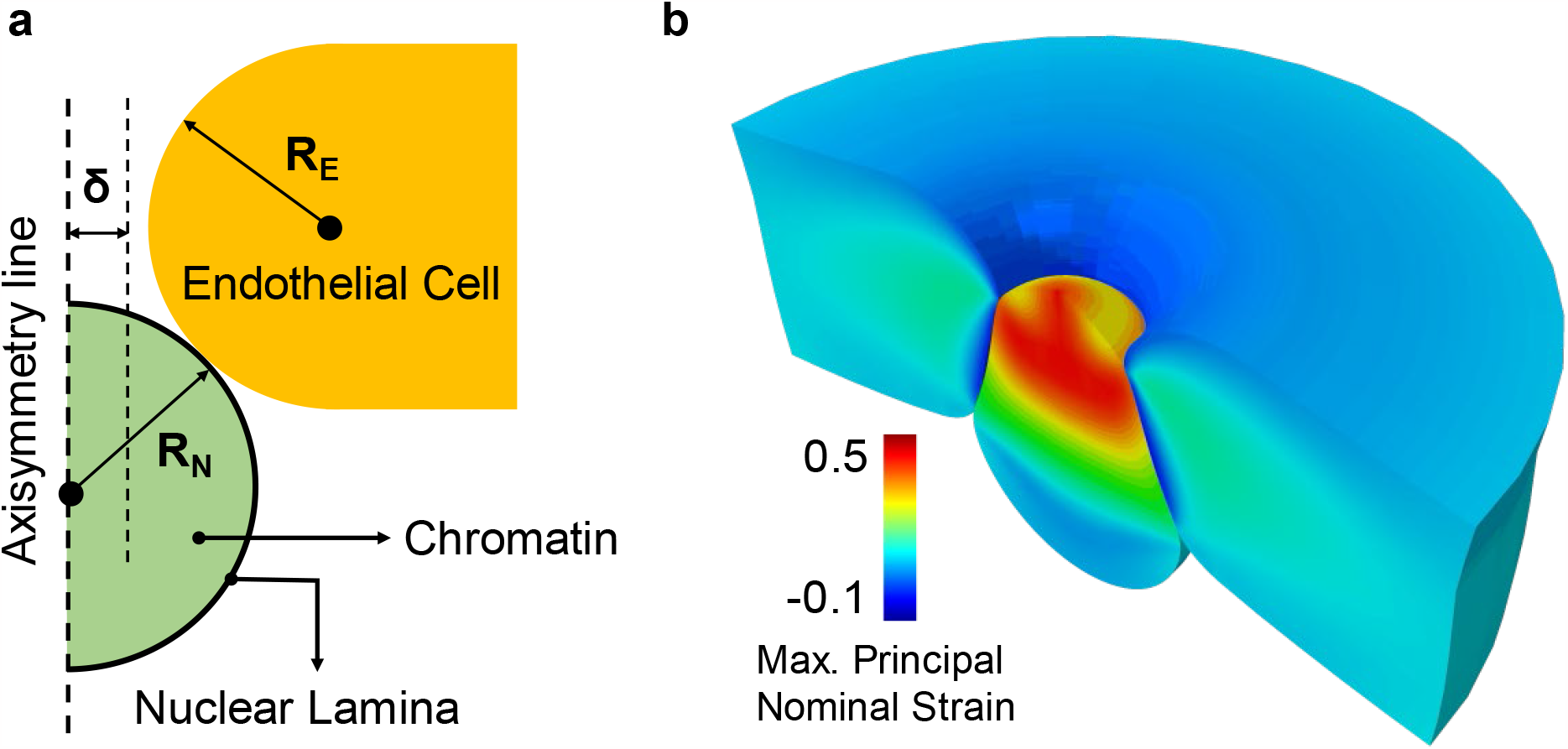
Finite element modeling of nucleus transmigration through endothelial pore. (a) This figure shows a schematic of pre-deformation nucleus and endothelial cell. The radius of both nucleus and endothelial cell were fixed at *R*_*N*_ = *R*_*E*_ = 3 μm. The pore radius *δ*varies from 1 to 2.25 μm. The nuclear lamina is modeled as a shell structure that encloses the interior chromatin solid. The endothelial cell is modeled both as a rigid structure and a hyperelastic solid. (b) The figure showcases a post-deformation 3D cut of the nucleus passing through the endothelial pore, with maximum principal nominal strain contoured for both the nucleus and endothelial cell.

### 3.5. Variable Density

Fig. 5 illustrates the impact of varying the network density *ρ*_*0*_ on nucleus mechanics and the nuclear lamina membrane properties in rigid gap transmigration simulations. In Fig. 5a, it is evident that the critical force *F*_*c*_ required to pass through the endothelial pore increases as the pore radius *δ* tightens, however, the curve slope starts to sharpen as the stiffening mechanism of dimers comes into play at roughly *δ ≤* 1 μm. This may imply the limiting contribution of the nuclear lamina to cell transmigration through narrow spaces. On the other hand, elevation in lamin concentration within the nuclear lamina can also be regarded as an active response to restrict transmigration.

**Fig. 5.**
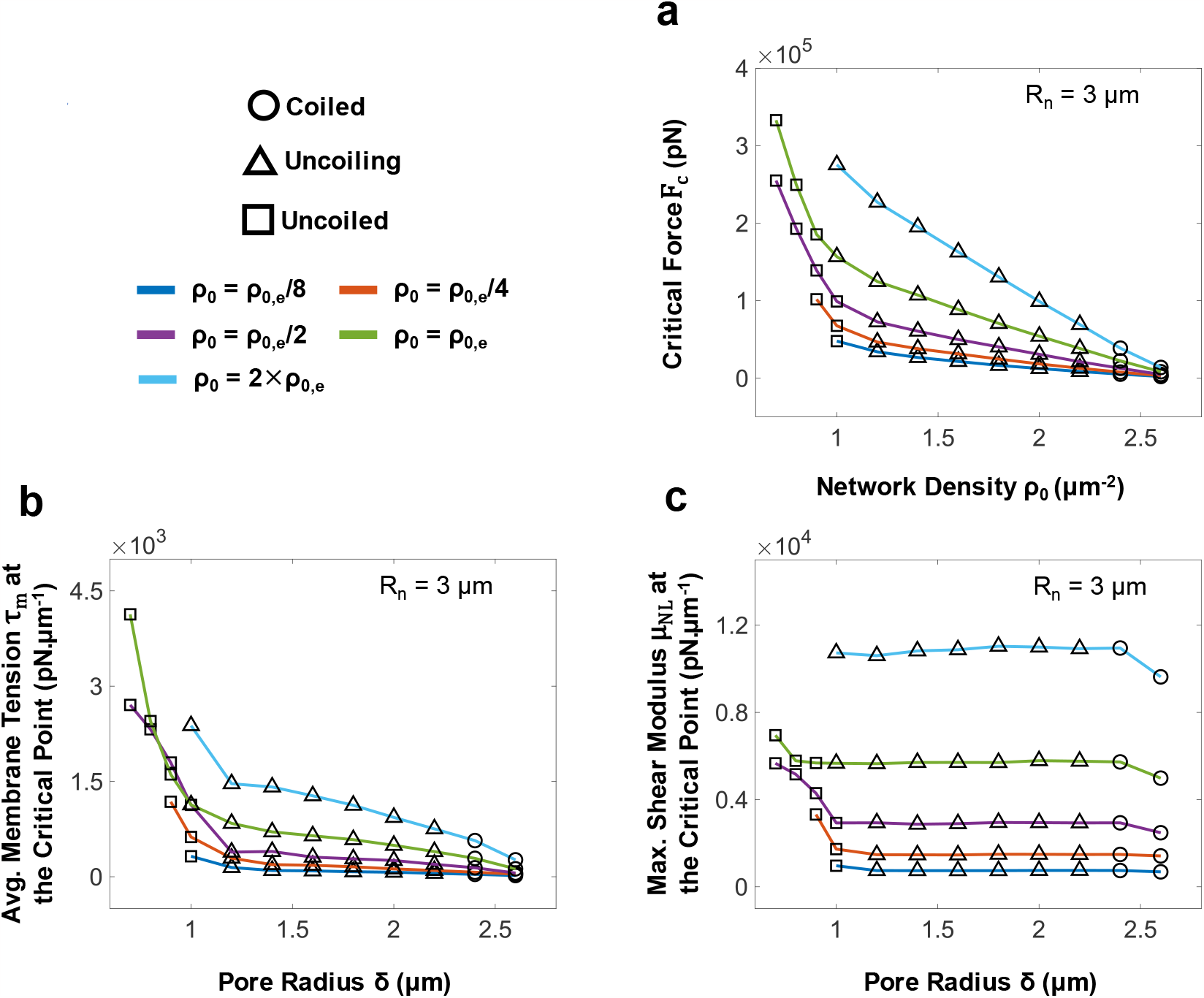
Effect of network density *ρ*_*0*_ in nucleus mechanics examined at the critical point of rigid gap transmigration finite element simulation. All other parameters are fixed at their fitting values. The three markers on top (circle, triangle and square) indicate the deformation regime of the filament with maximum extension. The nucleus radius is fixed at *R*_*n*_ = 3 μm and network densities employed for simulations are all ratios of *ρ*_*0,e*_. (a) The graph depicts the critical force necessary for the nucleus to squeeze *F*_*c*_, as a function of pore radius *δ*. When examining larger pores, the impact of density on transmigration becomes more discernible, but as the pore constricts, network density significantly affects nucleus mechanics. Furthermore, the sharp rise in critical force for *δ* smaller than 1μm underscores the macroscopic consequences of microstructural behavior. (b) The graph illustrates the average membrane tension *τ*_*m*_ within nuclear lamina shell elements, plotted against *δ*. As the density decreases, the nuclear lamina necessitates less tension to undergo the same strain, resulting in a reduction in the maximum membrane tension it experiences. The distinct extension phases of the lamin dimer are evident in the slope of membrane tension curve, with *δ* less than 1μm showing a sharp growth. (c) This graph displays the maximum shear modulus of the nuclear lamina network *μ*_*NL*_ at the critical point of transmigration, plotted against *δ*. Similar to membrane tension, the shear modulus during nucleus transmigration is highly dependent on the density of the nuclear lamina. There is a nearly constant maximum shear modulus with respect to the pore size given certain network density, which lies within a stability buffer at very large pore radii, and the final stiffening phase of constituent lamin filaments. This figure provides clear evidence that our model predicts the capacity of nuclear lamina to support nucleus transmigration while also maintaining the shape of nucleus by absorbing minor external forces, as well as protecting the chromatin from damaging strains.

In Fig. 5b, the average membrane tension *τ*_*m*_ achieved at the critical point (associated with emergence of critical force) is recorded and graphed for variable *ρ*_*0*_ and *δ*. Microstructural mechanics of lamin dimers are evidently reflected in the membrane tension, for distinct uncoiling and stiffening behaviors are produced for *δ* less than and more than 1 μm, respectively. As indicated by the trend in *F*_*c*_, membrane tension curves are in direct influence of the network density.

Fig. 5c illustrates that alterations in network density have a notable impact on the maximum shear modulus (*μ*_*NL*_) at the critical point across the entire range of pore radii, maintaining a stable value. However, the stable shear modulus levels, given a certain network density *ρ*_*0*_, is encompassed within an initial rise to that level and a subsequent increase to larger shear moduli. As elaborated in section 3.2, the change in shear modulus due to the incorporation of the BT model in the microstructure-based formulation highlights the role of the nuclear lamina as a stabilizing buffer, preserving the nucleus’s stability in the face of minor mechanical fluctuations and coincidental stresses.

Research indicates that under high curvature conditions, the lamin B network undergoes a dilution process to accommodate and facilitate greater deformations [20]. During transmigration, the change in the network density of the lamin can act as a crucial trigger, influencing whether the nucleus aims to prohibit or facilitate the process, therefore, the network density parameter in our microstructure-based model can serve as a valuable modeling parameter in cell mechanics simulations.

### 3.6. Variable *F*_*1/2*_ and *ΔΔx*^*\**^

Fig. 6a, Fig. 6c, and Fig. 6e demonstrate how the reduction in the uncoiling force (*F*_*1/2*_) can facilitate the transmigration of the nucleus. Similarly, Fig. 6b, Fig. 6d, and Fig. 6f show that reducing the uncoiling length (*ΔΔx*^*\**^) has a similar effect on transmigration, although reducing this parameter by the same factor as *F*_*1/2*_ results in a smaller change in the mechanics of the nucleus. Studies have revealed that tetrameric interactions, such as A11 and eA22, play a crucial role in stabilizing the coiled-coil domains in the dimer. Mutations in these interactions, observed in certain laminopathies, can lead to the destabilization of the lamin dimer, resulting in either softening or stiffening effects [46]. By considering the uncoiling parameters, *F*_*1/2*_ and *ΔΔx*^*\**^, the impact of these mutations on nucleus-related phenomena can be studied through finite element simulations.

**Fig. 6.**
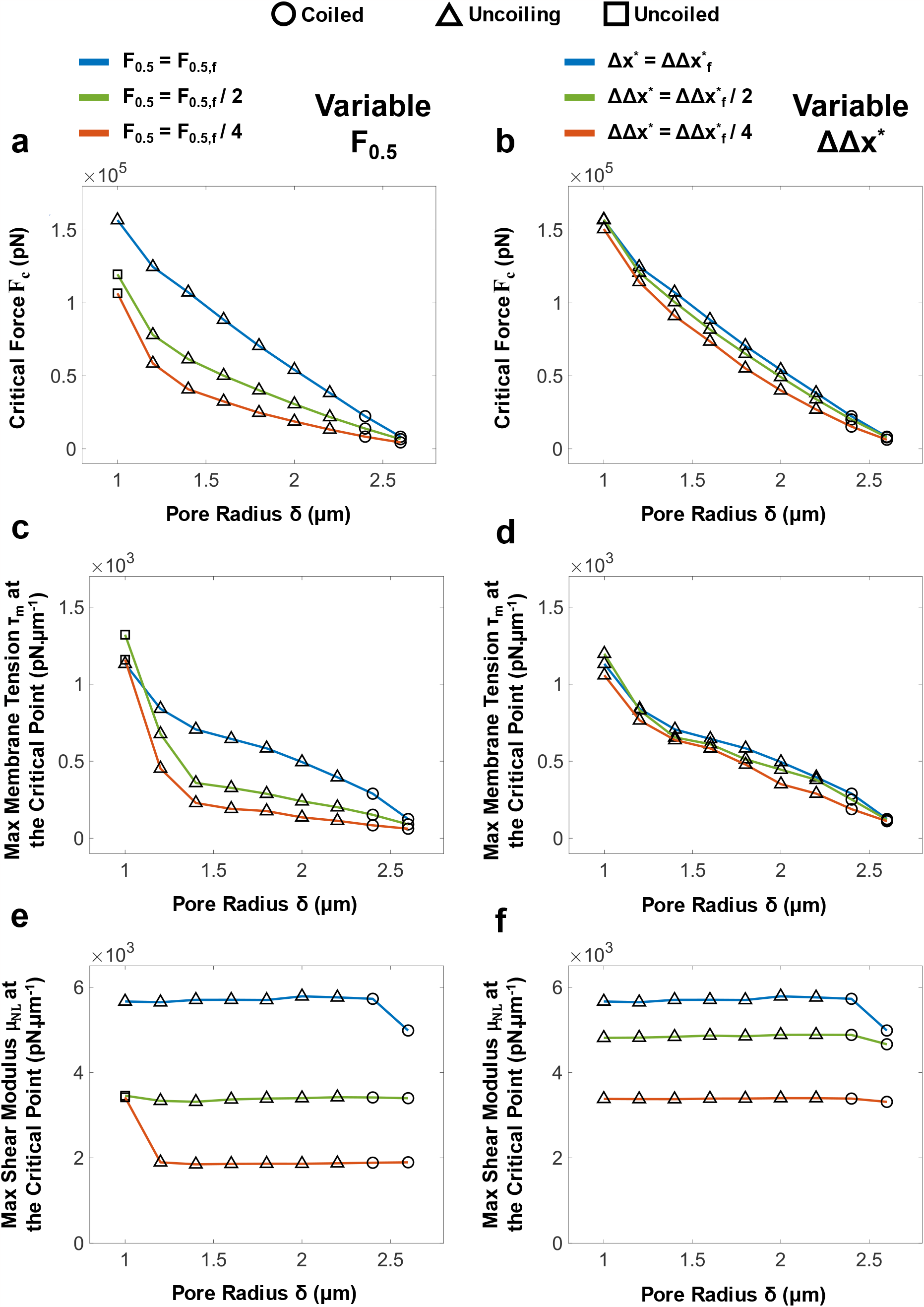
The effect of lamin dimer uncoiling factors *F*_*1/2*_ and *ΔΔx*^*\**^ in nucleus mechanics examined at the critical point of rigid gap transmigration finite element simulation. All other parameters are fixed at their fitting values, the nucleus radius is fixed at *R*_*n*_ = 3 μm and density is set at *ρ*_*0,e*_. Figures (a) and (b) display the critical transmigration force (*F*_*c*_) plotted against pore radius, with one factor reduced at a time: (a) shows the effect of reducing *F*_*1/2*_, and (b) represents the effect of reducing *ΔΔx*^*\**^. Although both factors lead to a decrease in the critical force, compared to uncoiling length *ΔΔx*^*\**^, reducing the uncoiling force (*F*_*1/2*_) has a more significant effect in facilitating the transmigration process. The impact of filament stiffening is evident in both cases, particularly for smaller values of *F*_*1/2*_, where uncoiling occurs at lower strains. Figures (c) and (d) depict the maximum membrane tension at the critical point plotted against pore radius, considering the same parameter changes as shown in (a) and (b), respectively. The maximum membrane tension decreases significantly more when reducing *F*_*1/2*_ compared to reducing *ΔΔx*^*\**^. The mechanical behavior of filaments is clearly reflected in the tension over the nuclear lamina. The Figure (e) and (f) demonstrate the trend of maximum shear modulus as a function of pore size, considering the same parameter changes as shown in (a) and (b), respectively. Reducing *F*_*1/2*_ has a more pronounced impact, leading to more frequent uncoiling and consequent disappearance of the nuclear lamina mechanical contribution on nucleus stiffness.

## 4. Discussion and Conclusion

We utilized a lamin A dimer structure based on X-ray diffraction crystallography to conduct a molecular dynamics simulation of protein stretching which provided us with the force-extension response of the dimer. We characterized deformation phases of the lamin A dimer and employed a physics-based semiflexible model to describe the pre-uncoiling mechanical behavior. For the post-uncoiling response, we fitted the data to a flexible WLC interpolation model. To model the uncoiling process, we incorporated a two-state kinetic transition fraction and combined Eq. 1 and Eq. 2 to create a comprehensive coiled-coil equation, describing the extension as a function of the pulling force. After developing expressions for the stress resultants of a 2D membrane containing lamin filaments, we derived equations for the membrane tension and the shear modulus of the nuclear lamina as a function of coiled-coil model parameters, filament characteristic orientations and network density. We approximated the initial shear modulus of the network and estimated the network density using available Cryo-ET image analysis data. Subsequently, we investigated the impact of the number of characteristic orientations on the mechanics of the nuclear lamina. Finally, we constructed a finite element model to simulate nucleus transmigration through an endothelial pore. In this model, we examined the influence of varying network density, uncoiling force, and uncoiling length across a range of pore radii on the mechanics of the nuclear lamina and the minimum force required for a successful transmigration.

Our model comprehensively integrates the microstructure and network properties of the nuclear lamina, which is the most critical mechanical component of the nucleus and helps us to study nucleus-related phenomena in greater detail. Incorporating BT model introduced a stability buffer in the definition of shear modulus which postulates the possible role of nuclear lamina in maintaining the shape of the nucleus under various micro-tensions. Also, the uncoiling mechanism introduces a permissible deformability limit for the nucleus in order to protect it during high-deformation incidents, asserting that nuclear lamina acts as a shock absorber for the nucleus [47].

In the future, this model holds the potential to enhance our understanding of the interplay between chromatin and the nuclear lamina in nucleus mechanics. Additionally, it can be used to investigate the effects of various nucleus abnormalities, such as the distinct mechanical response of progerin compared to lamin dimer and the upregulation or downregulation of lamin in different cancers, on cell mechanical function.

